# Genetic and antigenic characterization of an expanding H3 influenza A virus clade in US swine visualized by Nextstrain

**DOI:** 10.1101/2021.11.17.469008

**Authors:** Megan N. Neveau, Michael A. Zeller, Bryan S. Kaplan, Carine K. Souza, Phillip C. Gauger, Amy L. Vincent, Tavis K. Anderson

## Abstract

Defining factors that influence spatial and temporal patterns of influenza A virus (IAV) is essential to inform vaccine strain selection and strategies to reduce the spread of potentially zoonotic swine-origin IAV. The relative frequency of detection of the H3 phylogenetic clade 1990.4.a (colloquially known as C-IVA) in US swine declined to 7% in 2017, but increased to 32% in 2019. We conducted phylogenetic and phenotypic analyses to determine putative mechanisms associated with increased detection. We created an implementation of Nextstrain to visualize the emergence, spatial spread, and genetic evolution of H3 IAV-S, identifying two C-IVA clades that emerged in 2017 and cocirculated in multiple US states. Phylodynamic analysis of the HA gene documented low relative genetic diversity from 2017 to 2019, suggesting clonal expansion. The major H3 C-IVA clade contained an N156H amino acid substitution, but HI assays demonstrated no significant antigenic drift. The minor HA clade was paired with the NA clade N2-2002B prior to 2016, but acquired and maintained N2-2002A in 2016, resulting in a loss in antigenic cross-reactivity between N2-2002B and −2002A containing H3N2 strains. The major C-IVA clade viruses acquired a nucleoprotein (NP) of the H1N1pdm09 lineage through reassortment in replacement of the North American swine lineage NP. Instead of genetic or antigenic diversity within the C-IVA HA, our data suggest that population immunity to H3 2010.1, along with antigenic diversity of the NA and acquisition of the H1N1pdm09 NP gene likely explain the re-emergence and transmission of C-IVA H3N2 in swine.

**Importance:** Genetically distinct clades of influenza A virus (IAV) in swine undermines efforts to control the disease. Swine producers commonly use vaccines, and vaccine strains are selected by identifying the most common hemagglutinin (HA) gene from viruses detected in a farm or a region. In 2019, we identified an increase in detection frequency of an H3 phylogenetic clade, C-IVA, which was previously circulating at much lower levels in U.S. swine. Our study identified genetic and antigenic factors contributing to its resurgence by linking comprehensive phylodynamic analyses with empirical wet-lab experiments and visualized these evolutionary analyses in a Nextstrain implementation. The contemporary C-IVA HA genes did not demonstrate an increase in genetic diversity nor significant antigenic changes. N2 genes did demonstrate antigenic diversity, and the expanding C-IVA clade acquired a nucleoprotein (NP) gene segment via reassortment. Virus phenotype and vaccination targeting prior dominant HA clades likely contributed to the clade’s success.

## Introduction

Influenza A virus (IAV) is an economically important pathogen of swine that has the ability to evolve and evade the host immune response which presents a challenge to current control strategies. The negative-sense, single-stranded RNA genome consists of eight non-contiguous gene segments that are known to encode between 10 and 17 proteins (1–3). The segmented genome structure creates opportunity for reassortment when two or more IAV strains concurrently infect the same host, resulting in novel gene combinations and increased diversity at the genomic level (4, 5). IAV can also rapidly accumulate mutations due to errors introduced by viral polymerase (6). Swine have both α2,6- and α2,3-Gal-linked sialic acid on the surface of their respiratory epithelial cells; therefore, in addition to swine-to-swine transmission, they have potential for infection from human and avian origin IAVs (7, 8). Consequently, observed IAV diversity in swine is increased by the transmission, occasional establishment, and evolution of avian and human IAV in swine populations.

The genetic diversity of IAV translates to a similarly large breadth of antigenic diversity in the hemagglutinin (HA) and neuraminidase (NA) proteins. The accumulation of amino acid substitutions from polymerase mutation in the surface glycoproteins often results in changes in the antigenic phenotype of IAV (9, 10). For the H3 subtype, a small number of amino acid residues have a disproportionate effect on antigenic phenotype in both humans and swine (11–13). Six of these amino acid positions (145, 155, 156, 158, 159, and 189; H3 mature peptide numbering (14)) are referred to as the H3 antigenic motif in swine (15). These six residues are located on the globular head of the HA protein and are adjacent to the receptor binding site. There may be constraints on substitution flexibility at these positions due to necessary conservation of receptor binding functionality, as shown for position 145 (13). Minimal genetic change may result in significant antigenic change, reducing the efficacy of current IAV vaccines against clinical disease and transmission in swine (16). Vaccination with whole inactivated virus (WIV) with oil in water adjuvant is common in swine in the United States (U.S.). However, WIV are most efficacious when the vaccine and challenge strains are closely related (17, 18). Thus, an important utility of IAV surveillance and sequence data in swine populations is to inform vaccine strain selection.

In 1998, investigations into severe respiratory disease in swine in the U.S. revealed the introduction of the H3N2 subtype of IAV into swine. The H3N2 that persisted had triple reassortant internal genes (TRIG) with HA, NA, and PB1 gene segments derived from human seasonal H3N2; PB2 and PA gene segments from avian IAV; and NP, M, and NS gene segments from classical swine H1N1 (19–21). The HA gene from this introduction established the colloquially named H3 cluster IV (C-IV) in the U.S. (named the 1990.4 lineage in global H3 nomenclature (21)). The C-IV HA continued to circulate in the U.S. and evolved into genetically distinct clades A through F (22). The C-IV clade A (C-IVA) began to increase in detection frequency beginning in 2010 and was the predominant H3 clade in US swine until 2016, when it was surpassed by H3 2010.1, a more recent human-seasonal incursion that subsequently became established as a swine lineage (21, 23, 24).

In this study, we detected a resurgence in C-IVA sequence detections in 2019, as well as a relative decrease in detection of H3 2010.1. We quantified genetic and antigenic characteristics associated with the increased detection frequency of the C-IVA clade. Concurrently, we adapted the Nextstrain platform (25) to IAV in swine to provide near real-time phylogenetic visualization of surveillance data for the H3 subtype. Collectively, these analyses provide insight into the factors contributing to the expansion of swine IAV clades and improve our ability to predict mechanisms that allow IAV to evade current control measures in swine populations.

## Results

### Increased detection frequency followed by increased relative genetic diversity of H3 Clade IV-A

From 2011 to 2015 the C-IVA genetic clade was detected more frequently in U.S. swine than the 2010.1 genetic clade (Figure 1a). Between 2011 and 2014, the C-IVA clade paired with a N2 2002B gene was the primary HA-NA pairing detected among H3N2, while the C-IVA clade paired with a N2 2002A gene had decreased detection frequency. From 2015-2017, the C-IVA clade paired with a N2 2002B gene decreased in detection frequency, and detections of the 2010.1 HA clade paired with N2 2002A and 2002B genes increased from less than 20% to more than 90%. The majority of 2010.1 viruses detected were paired with a N2 2002B. In 2018, the H3 clade frequency patterns abruptly changed again, with the C-IVA at 39% of H3 HA genes. In 2019, the C-IVA HA genes were detected at 75%, and in 2020 the C-IVA HA genes represented more than 80% of the H3 genes in surveillance. The patterns in detections of the C-IVA and 2010.1 H3 clades in the first three months of 2021 remained similar to that of 2020.

**Figure 1.**
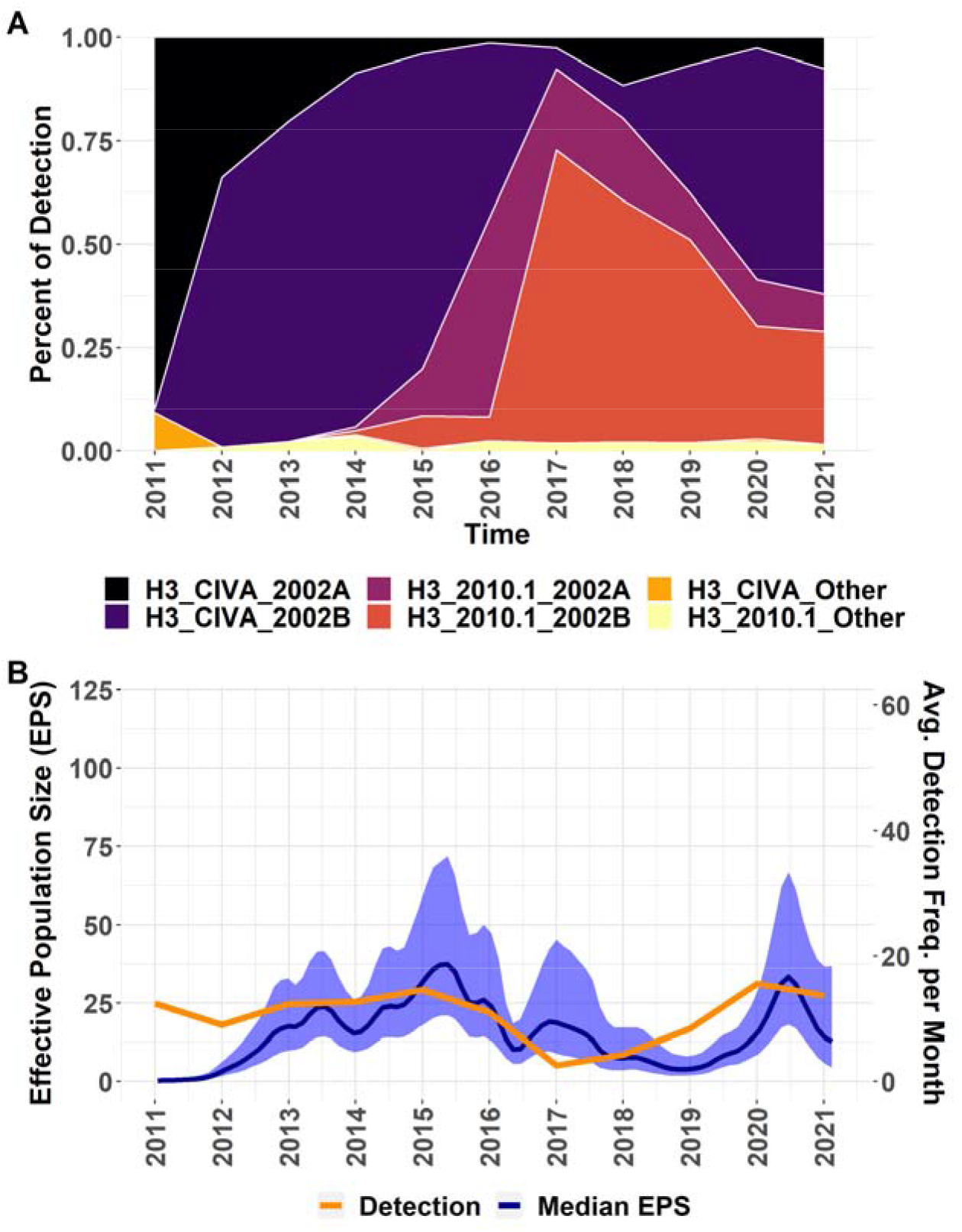
Relative C-IVA and 2010.1 HA-NA pair detection frequency and relative C-IVA genetic diversity from 2011 to 2021. **(A)** Proportional yearly detection frequency of H3 clades C-IVA and 2010.1 with the corresponding NA genetic clade from public data. In legend, HA clade is followed by NA clade. “Other” includes NA genes from the N2 1998 clade and one-off human-to-swine transmission events. **(B)** Effective population size (EPS) and average detection frequency per month of C-IVA viruses. EPS estimates relative genetic diversity within the HA genes of the C-IVA clade. Blue shading is 95% highest posterior density (HPD) interval.

To investigate the C-IVA expansion, we measured the median posterior rate of nucleotide substitution for the C-IVA clade by Bayesian analysis at 4.265 × 10^−3^ (95%HPD: 3.956 × 10^−3^, 4.589 × 10^−3^). Relative genetic diversity of the HA gene was estimated by a Bayesian demographic reconstruction and demonstrated an almost linear increase in median effective population size (EPS) in C-IVA genes from 2011 to 2015, but a decrease from 2015 to 2019 (Figure 1b). Despite minimal relative genetic diversity in 2018 and 2019, detection frequency began to increase. The increase in relative genetic diversity in 2020 lagged behind the increase in detection frequency. The trends in relative genetic diversity are supported by the topology of a maximum-likelihood phylogeny (Supplementary Figure 1) with external branches that are shorter relative to branches on the interior of the tree, i.e., the topology of the phylogeny is balanced. The equal and undirected propagation of leaves along the branches suggests that selection is not acting on the HA gene at the time of this study, as HA under selection would appear as a “ladder-like” topology of the gene tree.

### Two co-circulating clades with onward transmission after 2018

The HA tree (Figure 2) shows many co-circulating C-IVA genetic clades that corresponded with high levels of relative genetic diversity in 2013 and 2015 (Figure 1b). After 2018, only two distinct genetic clades of the C-IVA clade were apparent: a major clade representing 245 detections (71 in 2019, 174 in 2020) and a minor clade representing 24 detections (19 in 2019 and 6 in 2020). The major clade viruses were first detected in the Southwest region (Texas, Oklahoma, and Kansas) of the U.S. in 2017 and 2018, but were detected in the major pork producing states of the Midwest by January 2019. By late-2019, major clade viruses were also detected in North Carolina and some less hog-dense states such as Michigan and Pennsylvania. Although there was broad geographic representation, 56% of the detections of this clade were in Iowa and Indiana. The minor clade was initially detected in the Midwest with only 5 detections outside this region from 2018 to present.

**Figure 2.**
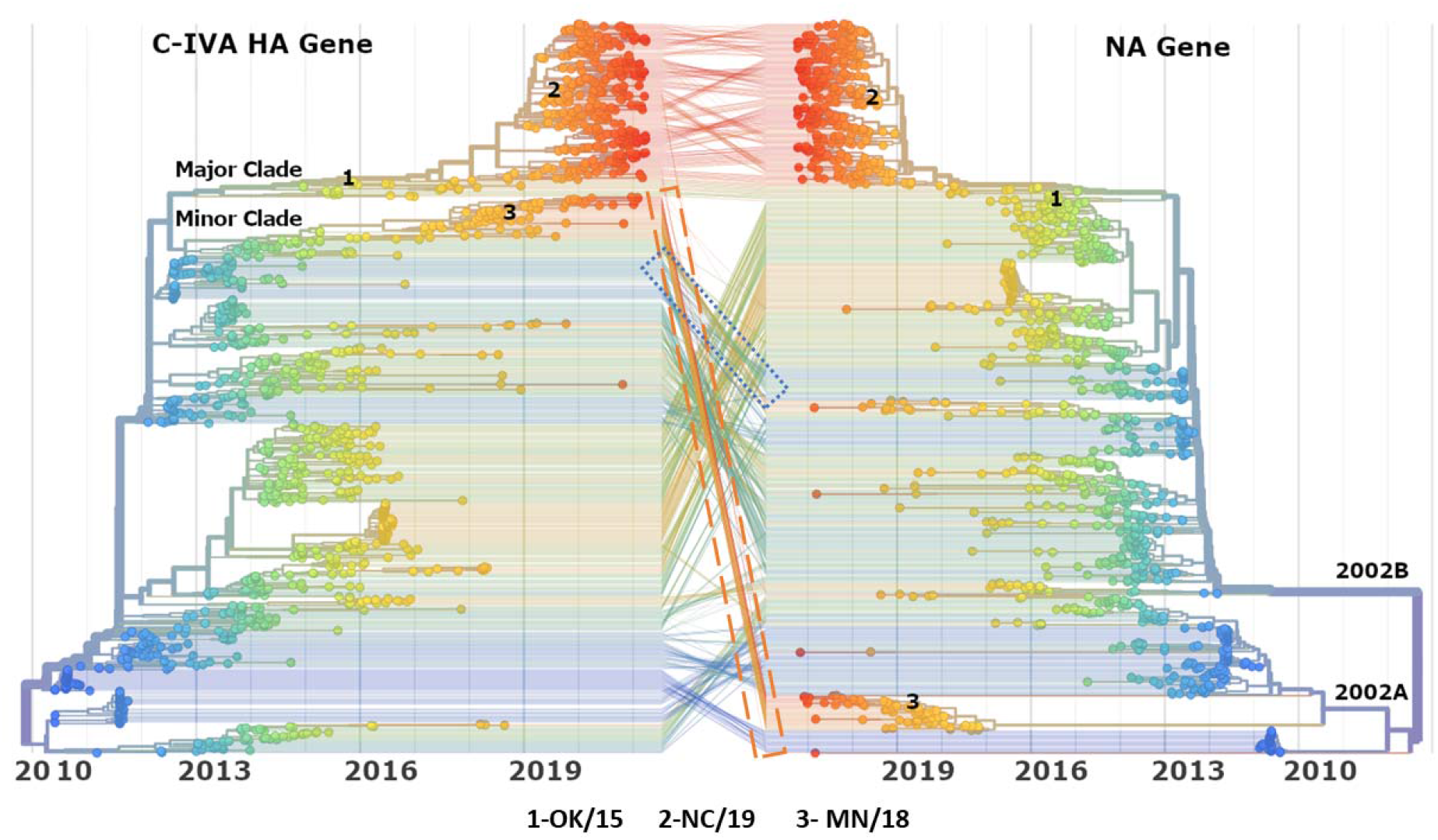
Tanglegram of corresponding C-IVA HA and NA gene time-scaled phylogenetic trees with sequences from 2010 to March 2021. The major and minor contemporary C-IVA clades are labeled on the HA tree. N2 genetic clades are labeled on the NA tree. The three contemporary C-IVA strains used in antigenic analysis are also labeled on both trees. (1: A/swine/Oklahoma/A01770191/2015, OK/15; 2: A/swine/North Carolina/A02245294/2019, NC/19; and 3: A/swine/Minnesota/A02266068/2018, MN/18). Lines between the two trees indicate HA-NA pairing. Tree branches and leaves colored on blue to red gradient by yearly progression of time in accordance with the x-axis. Blue, dotted line indicates pairing of the minor clade with N2 2002B prior to reassortment and orange, dashed line indicates reassortment of the minor clade with N2 2002A. The Nextstrain platform can be used to visualize the tanglegram in finer detail at https://flu-crew.org/

### Reassortment with the H3 IV-A and novel NA and NP gene segment pairings

The HA gene segments of the major C-IVA clade paired consistently with N2 clade 2002B.2 gene segments, matching the topology of the congruent NA phylogenetic tree (Figure 2). The ancestral viruses of the minor clade paired with N2 2002B.2 from mid-2012 to 2016. In 2016 (95% CI: 2016-06-05, 2017-02-08), the minor clade showed evidence of reassortment with a genetically distinct N2 2002A.2 clade that no other C-IVA HA gene segment was paired with in the past decade. To observe patterns in the available genomes of the IVA H3N2 strains, the lineages of the remaining six gene segments were annotated onto the HA tree (Figure 3). Shortly after the 2009 H1N1 pandemic, North American swine IAV acquired the M gene from the H1N1pdm09 lineage, but retained a nucleoprotein (NP) gene segment from the triple-reassortment H3N2 (TRIG) lineage. The major clade showed evidence of reassortment with an NP from the H1N1pdm09 lineage beginning in 2017. All available WGS (n=58) from the major clade contained a pdm09 lineage NP after September 2018. There was no evidence of replacement of the TRIG lineage in the PB2, PB1, PA or NS gene segments.

**Figure 3.**
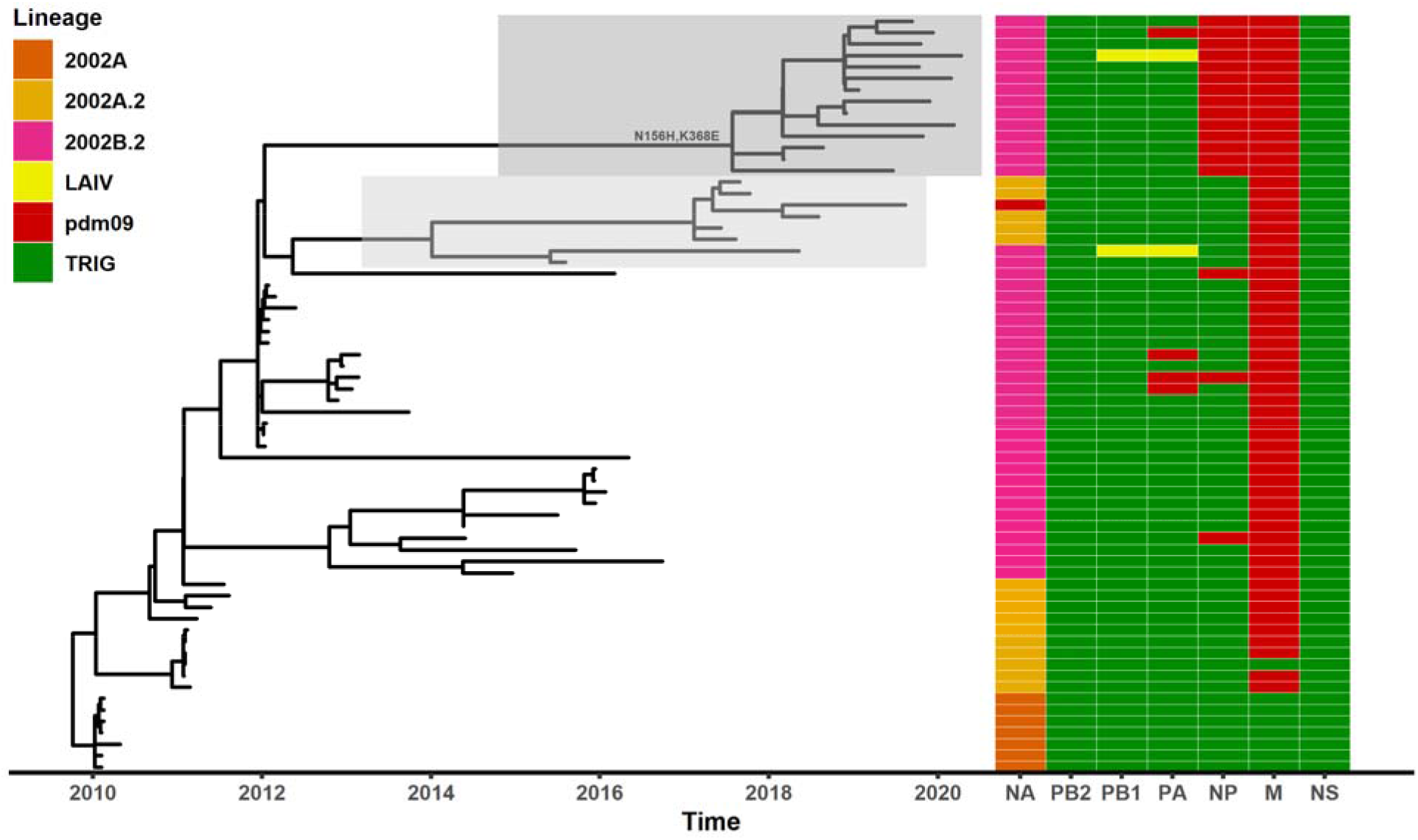
C-IVA HA time-scaled tree with genome constellation. The corresponding NA and internal gene (PB2, PB1, PA, NP, M, NS) lineage of each tip are indicated by the color-coded rectangles on the right side of figure. The major contemporary C-IVA clade is boxed in dark grey on the tree and the minor contemporary C-IVA clade is boxed in light grey on the tree. The two substitutions defining the initial expansion of the major clade (N156H and K368E) are annotated onto the tree. Dark Orange=N2 2002A; Light Orange = N2 2002A.2; Pink = N2 2002B.2; Yellow=LAIV; Red=H1N1pdm09 lineage; Green=swine triple reassortant internal gene (TRIG) lineage.

### N156H antigenic motif substitution

The time-scaled tree annotated with amino acid substitutions created with Nextstrain showed two amino acid substitutions associated with the expansion of the major clade, N156H and K368E, dating to November 2016 (95% CI: 2016-06-04,2017-05-18) (Figure 3). The 156 position was identified in the previously characterized antigenic motif (11, 12, 15). Thirteen other amino acid substitutions were detected in H3 genes in the major clade after December 2016 but with no consistent pattern. Five of the thirteen occurred in antigenic regions of HA1 (N96S, S146G, N158D, I214V, V323I; H3 mature peptide numbering) (26).

### Antigenic diversity of HA

Contemporary strains were selected to represent three groups of C-IVA for antigenic assessment of the HA: the major clade prior to the N156H substitution (A/swine/Oklahoma/A01770191/2015; OK/15; annotation 1 in Figure 2) with a motif of NYNNYK; the major clade following the N156H substitution (A/swine/North Carolina/A02245294/2019; NC/19; annotation 2 in Figure 2) with a motif of NYHNYK; and the minor clade (A/swine/Minnesota/A02266068/2018; MN/18; annotation 3 in Figure 2) with a motif of NYNNYK. Significant antigenic drift is defined by an 8-fold loss in HI cross-reactivity, which corresponds to 3 antigenic units (AU) between viruses in an antigenic map. The three contemporary C-IVA representative strains were within 2 AU of each other and within 3 AU of the other C-IVA tested strains (Figure 4a). Two reference antigens from a different genetic clade, C-IVB, were near the 3 AU significance cutoff. The C-IVB strain representing the NYHNYK antigenic motif was closer to all three tested C-IVA viruses, particularly NC/19 which had the same NYHNYK antigenic motif, than the CIV-B strain representing the NYNNYK antigenic motif. The representative strain of CIV-C showed significant antigenic distance of over 5 AU from the C-IVA strains.

**Figure 4.**
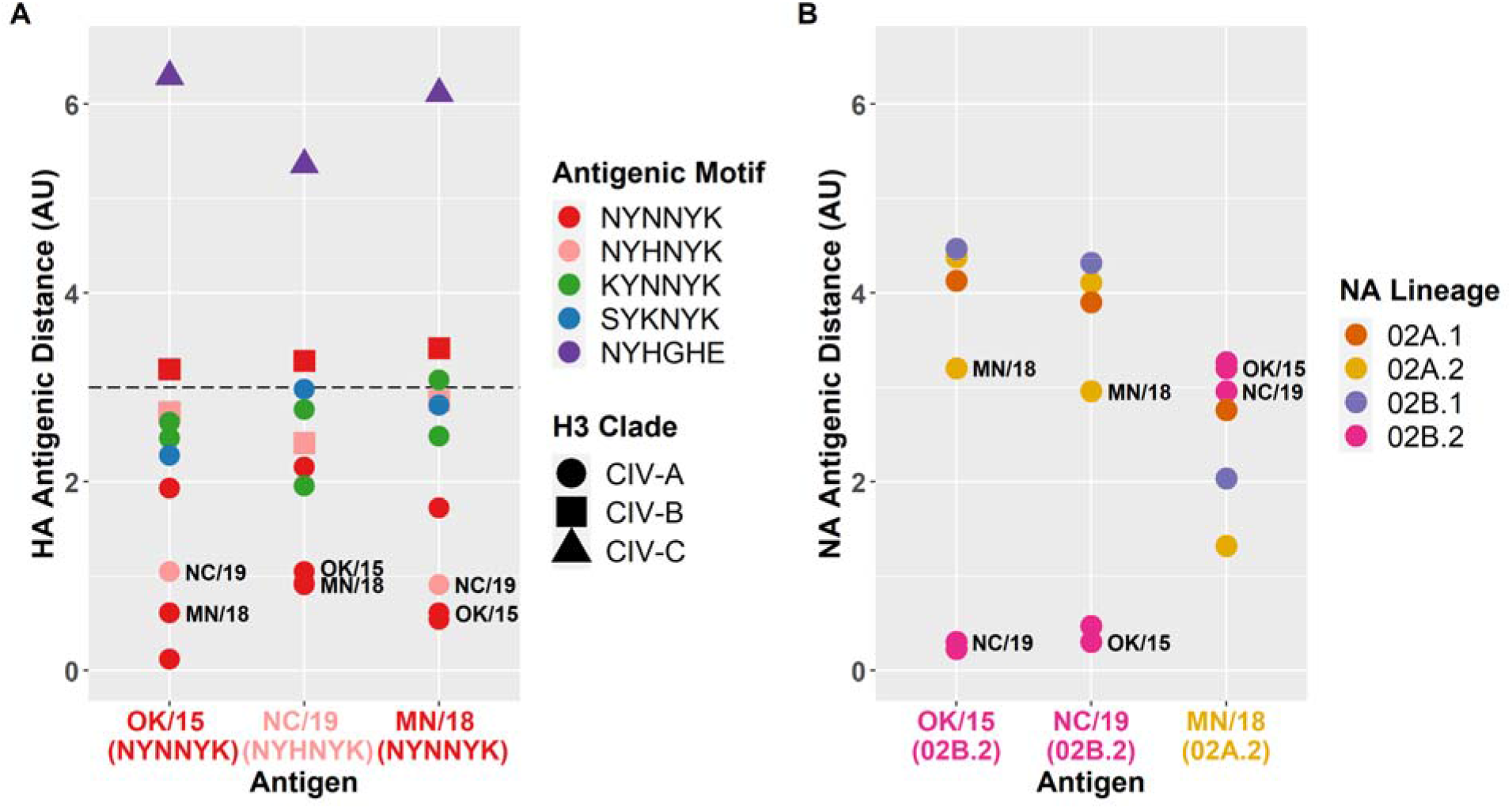
HA and NA antigenic distance. **(A)** HA antigenic distance between three contemporary C-IVA test antigens and relevant H3 reference antigens. Distances are computed by merging the raw HI results from the assay described in this experiment with results from previous HI assays in ACMACS. Points are colored by H3 antigenic motif and their shape corresponds to their H3 clade classification. The three test antigens on the x-axis are colored by H3 antigenic motif. Significant HA antigenic drift is defined as antigenic distance of at least 3 AU and is denoted by the black dashed line. **(B)** NA antigenic distance between three contemporary C-IVA test antigens and antigens from four N2 2002 lineages. Distances are computed by merging the raw NI results from the assay described in this experiment with results from previous NI assays in ACMACS. Points are colored by NA lineage. The three test antigens on the x-axis are also colored by NA lineage. The test antigens, OK/15, NC/10 and MN/18, are labeled with black text in both panels.

### Neuraminidase Inhibition and diversity with Enzyme-Linked Lectin Assay (ELLA)

The NI titers of the same three HI test strains were assessed against reference antisera of swine N2 clades: the OK/15 and NC/19 strains had a N2-2002B.2 gene; the minor clade representative MN/18 had a N2-2002A.2 gene. These new NI titer data were combined with previous experiments from Kaplan et al. (27), and antigenic distances were extracted. There were differences in NI antigenic phenotype within the N2-2002 lineage, which was further divided into 2002A.1, 2002A.2, 2002B.1 and 2002B.2 (27, 28). In particular, the minor clade MN/18 virus with the 2002A.2 lineage had an antigenic distance of 3 AU from the 2002B.2 lineage viruses, including the major clade NC/19 virus (Figure 4b). Despite being isolated 4 years apart, the 2002B.2 lineage viruses retained NI antigenic properties and were within 0.5 AU of each other.

## Discussion

This study investigated the recent increase in detection of H3 C-IVA viruses in US swine and circulating H3 diversity within swine populations could help explain genetic clade turnover and dominance dynamics. The H3 2010.1 clade that emerged in 2012 began to supplant the C-IVA clade in 2016 (Figure 1a). There was limited serologic cross-reactivity between 2010.1 and C-IVA swine H3N2 viruses (24). Animals in the population at ∼2016 were more likely to have natural immunity against 2010.1 viruses, while animals with natural immunity against C-IVA viruses gradually decreased. In addition, many herds were vaccinated against the 2010.1 clade of viruses with custom or autogenous vaccines following its emergence and dominance over C-IVA. Waning immunity against C-IVA viruses due to decreased natural immunity and increased focus on vaccines containing 2010.1 viruses could have provided an advantage to the C-IVA viruses. This is supported by the balanced shape of the HA phylogeny which suggests an absence of immune or vaccine driven selection within the emerging C-IVA clades (Figure 2, Figure 3).

Though the number of C-IVA detections increased since 2018, the relative genetic diversity within the clade did not increase until mid-2020. The 2018-2019 clonal expansion of C-IVA with low diversity suggests that a selective sweep occurred in the population. Sweep-related changes have been identified in human seasonal H3N2 IAV and most often detected at amino acid sites located on the HA (29, 30). In 2019, the relative genetic diversity began to rise, likely as the result of the success, spread, and subsequent diversification of the virus. The pattern of increasing detection frequency paired with low relative genetic diversity can act as an early warning signal that can be used to flag genetic clades of swine IAV that require characterization and risk assessment for swine agriculture and pandemic preparedness.

To determine whether a selective sweep occurred due to antigenic drift, we identified amino acid substitutions sustained in the major and minor clades that were circulating as detection frequency increased from 2018 to 2020. We identified an N156H substitution in the HA of the major clade. The 156 amino acid position was previously identified as having an effect on the antigenic phenotype, usually in combination with substitutions in other positions on the HA (11, 15, 31), but the impact of an amino acid substitution depends on the biological properties of the specific amino acid(s) that have changed in the context of the overall HA1 amino acid sequence (32, 33). This N156H substitution prompted the antigenic characterization of the major and minor clades via HI assays to assess for a potential loss in cross-reactivity. A significant loss in cross-reactivity of the contemporary 156H from prior strains with 156N would suggest a potential lack of population immunity that could explain the increased frequency of the major clade. However, our data did not demonstrate that the substitution caused significant antigenic drift. Antibodies raised against ancestral C-IVA demonstrated HI cross-reactivity against the more recent strains regardless of the substitution. Our results support previous findings that variation at position 156 alone did not cause significant antigenic drift (15, 31). The limited change in antigenic phenotype suggests the N156H substitution may not have been the primary cause of the observed clonal expansion of the C-IVA major clade.

With no evidence of significant antigenic drift in the HA between the contemporary C-IVA major and minor clades, the viruses were analyzed for evidence of other genetic signatures associated with the expansion. The minor clade recently reassorted to acquire N2-2002A.2 genes, while the major clade remained paired with N2-2002B.2. We further investigated the antigenic effects of this reassortment event with a panel of NI anti-sera previously used to describe antigenic variation among and between swine N2 lineages (27). The 2002B.2 of the C-IVA major clade viruses retained close antigenic relationships to the 2002B.2 reference antigen, but antigenic variation existed between clade representatives within the 2002 lineage. Since the early emerging 2010.1 viruses were paired with 2002A at the time they began to outnumber the prior circulating C-IVA, population immunity may have been skewed toward both a mismatched HA and NA. The 2002B.2 clade of N2 now represents the majority of circulating N2 detected in the swine population, so the importance of NA immunity to the maintenance of the major C-IVA H3 clade paired with 2002B is not clear.

We analyzed WGS data for evidence of reassortment of the internal genes. The major C-IVA clade was determined to have reassorted to acquire an NP of the H1N1pdm09 lineage. The genotype of IAV-S internal genes was summarized as a concatenation of one-letter codes representing the genetic lineage of each gene segment (PB2, PB1, PA, NP, M, and NS) without the HA and NA segments. The C-IVA internal gene constellation in 2010 was TTTTTT, with all internal genes coming from the TRIG lineage. C-IVA viruses then acquired a matrix gene from the H1N1pdm09 virus, with constellation TTTTPT, and this was common between 2009 and 2016 (34). This constellation was also involved in an H3N2v outbreak in humans in 2011-2012, causing 340 cases across 13 US states (35). However, beginning in 2017, the internal gene constellation found in the C-IVA major clade that increased in detection frequency was TTTPPT with a H1N1pdm09 NP gene. This constellation was observed before, but was uncommon (22 of the 368 isolates between 2009 and 2016) (34). The influenza NP is characterized as a structural RNA-binding protein that forms the ribonucleoprotein (RNP) particle (36), and its genetic variation may impact functions such as temporal regulation of apoptosis or import and export of vRNPs from the nucleus (37, 38). Results from a transmission study in pigs have demonstrated that H3 viruses with the TTTPPT constellation are more effective in viral transmission when compared H3 strains with a TTTTPT constellation (34). Consequently, our data suggest that success of this clade of viruses could be explained by differences between the pdm09 and TRIG genetic lineages of the NP acquired following reassortment in 2017.

Since H3 C-IVA viruses continue to make up roughly one-half of H3N2 detections within the national USDA influenza A virus in swine surveillance program in 2021, the increased detection frequency of C-IVA suggests that vaccines should include antigens from this clade of IAV. Subsequent surveillance is necessary to determine if vaccination against C-IVA will result in a decrease in detection; however, this would require additional knowledge of farm specific vaccines and vaccination strategies. Matching vaccine components to circulating diversity and understanding how swine transportation patterns and biosecurity practices affect the transmission of swine IAV H3 clades can help improve animal health. Further, the resurgence of this clade creates concern for public health, with the knowledge that a virus from the same clade was involved in causing a human outbreak in the context of reassortment. Our HI assay included a representative strain (A/swine/New York/A01104005/2011) that was genetically similar to the H3N2v from the 2011-2012 outbreaks and showed that the contemporary C-IVA representatives tested had not undergone significant antigenic drift. However, dominant swine H3N2 clades have caused numerous zoonotic events through human-swine agricultural interfaces (35, 39, 40), and more recent contemporary C-IVA swine strains may be antigenically drifted from the pandemic preparedness candidate vaccine virus A/Minnesota/11/2010 (41). Thus, understanding the factors that contribute to IAV in swine clade expansion is necessary to inform and improve prediction methods for more successful control measures, and to provide insight into pandemic preparedness efforts.

## Materials and Methods

### Data Collection

All available U.S. swine H3 nucleotide sequences (n=3395) detected between January 2010 and March 2021 deposited into GenBank (42) were downloaded from the Influenza Research Database (IRD) (43). Duplicate strains were removed from the dataset. Sequences were then classified using octoFLU (44), and those that were from the C-IVA clade (n=1376) were retained for further analysis. All available corresponding NA gene segments (n=1291) and whole genome sequences (WGS, n=545) were collated and classified into lineage using octoFLU (44) following the N2-1998 and N2-2002 clade divisions introduced by Kaplan et al. (27). H3 clade detection frequency was derived from public sequence data and validated with the private regional surveillance data housed in the Iowa State University Veterinary Diagnostic Lab (ISU VDL) visualized on ISU FLUture (23).

### Estimation of relative genetic diversity

To generate a computationally tractable dataset, we generated a random subset (n=500) of H3 C-IVA sequences via smof v2.21.0 (45). Sequences were aligned with mafft v7.450 (46) and a maximum-likelihood phylogenetic tree was inferred using the generalized time-reversible model (GTR) of nucleotide substitution in FastTree v2.1.11 (47). This tree was used in a root-to-tip regression analysis in TempEst v1.5.3 (48) to assess temporal signal and detect genes with incongruous genetic divergence and sampling dates. The final dataset (n=493) was then analyzed with BEAST v1.8.4 (49) to estimate effective population size of the C-IVA lineage over time. We applied the GMRF Bayesian Skyride coalescent model (50) with a GTR substitution model with gamma-distributed rate variation, an uncorrelated relaxed clock, and a MCMC chain length of 100,000,000 with sampling every 10,000 iterations. Demographic reconstruction was performed using the GMRF skyride reconstruction in Tracer v1.7.1 (51) and a maximum clade credibility (MCC) tree was generated using TreeAnnotator v1.8.4 (52).

### Deployment of Nextstrain for H3 IAV in swine

The Nextstrain (25) platform was adapted for H3 IAV-S. A time-scaled tree was estimated for all H3 swine IAV HA genes, and a focused C-IVA HA nucleotide sequence dataset using the “refine” Augur command (53). A separate time-scaled tree was estimated for paired NA nucleotide sequences and the two trees were then compared using the Auspice visualization platform (https://github.com/nextstrain/auspice). Amino acid substitutions were annotated on the backbone of the tree using the “ancestral” and “translate” commands. The H3 antigenic motif was visualized by combining the “Color By Genotype” function for positions 145, 155, 156, 158, 159, and 189. The lineages determined through octoFLU of the other six gene segments were mapped onto the HA tree using the “traits” command. The “traits” command also integrated geographic information at the U.S. state level and computed putative transmission between states. These data were exported as JSON files that are interactively visualized on the web at https://flu-crew.org/ on an AWS server using USDA-ARS SCInet with all files provided at https://github.com/flu-crew/h3-iva-evolution.

### Antigenic Characterization

We identified two C-IVA genetic clades co-circulating in the U.S. from January 2019 to Mar 2021; one clade formed the majority of detections (89.3% of 326 sequences) and the other was minor, but persistent. To identify a representative sequence for each clade, we generated an HA1 consensus sequence in Geneious Prime 2020.2.3 and selected the best matching field strain from the USDA IAV in swine surveillance system repository at the National Veterinary Services Laboratories, USDA-APHIS. For the major clade, we identified an amino acid substitution at position 156 on the backbone of the phylogeny using Nextstrain. We selected an “ancestral” strain and a “contemporary” strain to reflect the substitution at 156. Three field strains most similar to the consensus sequences were selected to be antigenically characterized: A/swine/Oklahoma/A01770191/2015 (OK/15) as the C-IVA major ancestral strain, A/swine/North Carolina/A02245294/2019 (NC/19) as the C-IVA major contemporary strain, and A/swine/Minnesota/A02266068/2018 (MN/18) as the C-IVA minor strain.

A panel of swine antisera was selected using sera previously produced by immunizing two pigs (11, 31). Antisera were treated with Receptor Destroying Enzyme (II) (Hardy Diagnostics), heat inactivated at 56°C for 30 min and adsorbed with 50% turkey red blood cells to remove nonspecific hemagglutination inhibitors. Hemagglutination inhibition (HI) assays were performed on test antigens and selected sera using turkey red blood cells (54) Cross-HI tables were merged and mapped in 3 dimensions following established antigenic cartography method (11). Antigenic distances between viruses were calculated in antigenic units (AU), in which 1 AU is equivalent to a 2-fold loss in HI cross-reactivity. A threshold of ≥3AU is considered significant loss in cross-reactivity (11). HI data with the selected H3 C-IVA strains were merged with a subset of previously generated H3 antigenic data and used to create three-dimensional antigenic maps using antigenic cartography (9, 11, 31). Antigenic distances between antigens generated in the 3-D map were extracted and plotted using ggplot2 in R (55).

An enzyme-linked lectin assay (ELLA) was used to determine neuraminidase inhibiting (NI) antibody titers. Serial dilutions of swine neuraminidase antisera were incubated with standardized concentrations of IAV in fetuin coated 96-well plates for 16 hrs at 37°C, 5% CO_2_. Cleavage of sialic acids from fetuin was detected using peanut agglutinin-horseradish peroxidase (PNA-HRP) (Sigma-Aldrich, St. Louis, MO) and 3,3’,5,5’-tetramethylbenzidine (TMB) (KPL Laboratories, Gaithersburg, MD), as previously described (56, 57). The optical density (OD) of the plates was read at 650 nm and the titer was assigned as the reciprocal of the highest dilution resulting in at least 50% inhibition.

### Data Availability

All sequence data are publicly available in NCBI Genbank and the code, data and supplemental materials associated with this manuscript provided at https://github.com/flu-crew/h3-iva-evolution. The Nextstrain for H3 swine IAV is hosted at https://flu-crew.org/.

## Acknowledgements

We gratefully acknowledge pork producers, swine veterinarians, and laboratories for participating in the USDA Influenza A Virus in Swine Surveillance System and publicly sharing sequences. This work was supported in part by: the Iowa State University Presidential Interdisciplinary Research Initiative; the Iowa State University Veterinary Diagnostic Laboratory; the U.S. Department of Agriculture (USDA) Agricultural Research Service [ARS project number 5030-32000-231-000-D]; the National Institute of Allergy and Infectious Diseases, National Institutes of Health, Department of Health and Human Services [contract number 75N93021C00015]; the USDA Agricultural Research Service Research Participation Program of the Oak Ridge Institute for Science and Education (ORISE) through an interagency agreement between the U.S. Department of Energy (DOE) and USDA Agricultural Research Service [contract number DE-AC05-06OR23100]; the Department of Defense, Defense Advanced Research Projects Agency, Preventing Emerging Pathogenic Threats program [contract number HR00112020034]; and the SCINet project of the USDA Agricultural Research Service [ARS project number 0500-00093-001-00-D]. The funders had no role in study design, data collection and interpretation, or the decision to submit the work for publication. Mention of trade names or commercial products in this article is solely for the purpose of providing specific information and does not imply recommendation or endorsement by the USDA, DOE, ORISE, DARPA, or ISU. USDA is an equal opportunity provider and employer.

## Supplementary Material

**Figure S1.** H3 C-IVA maximum likelihood tree. Branches are colored by the progression of time on a blue-to-red gradient beginning in 2010 and ending in March 2021.

**Table S1.** Results from a pair of hemagglutination inhibition (HI) assays performed on November 13, 2020 and January 22^nd^, 2021.

**Table S2.** Results from Neuraminidase Inhibition (NI) Assay.

