## Supplemental Material for "Genetic and antigenic characterization of an expanding H3 influenza A virus clade in US swine visualized by Nextstrain"

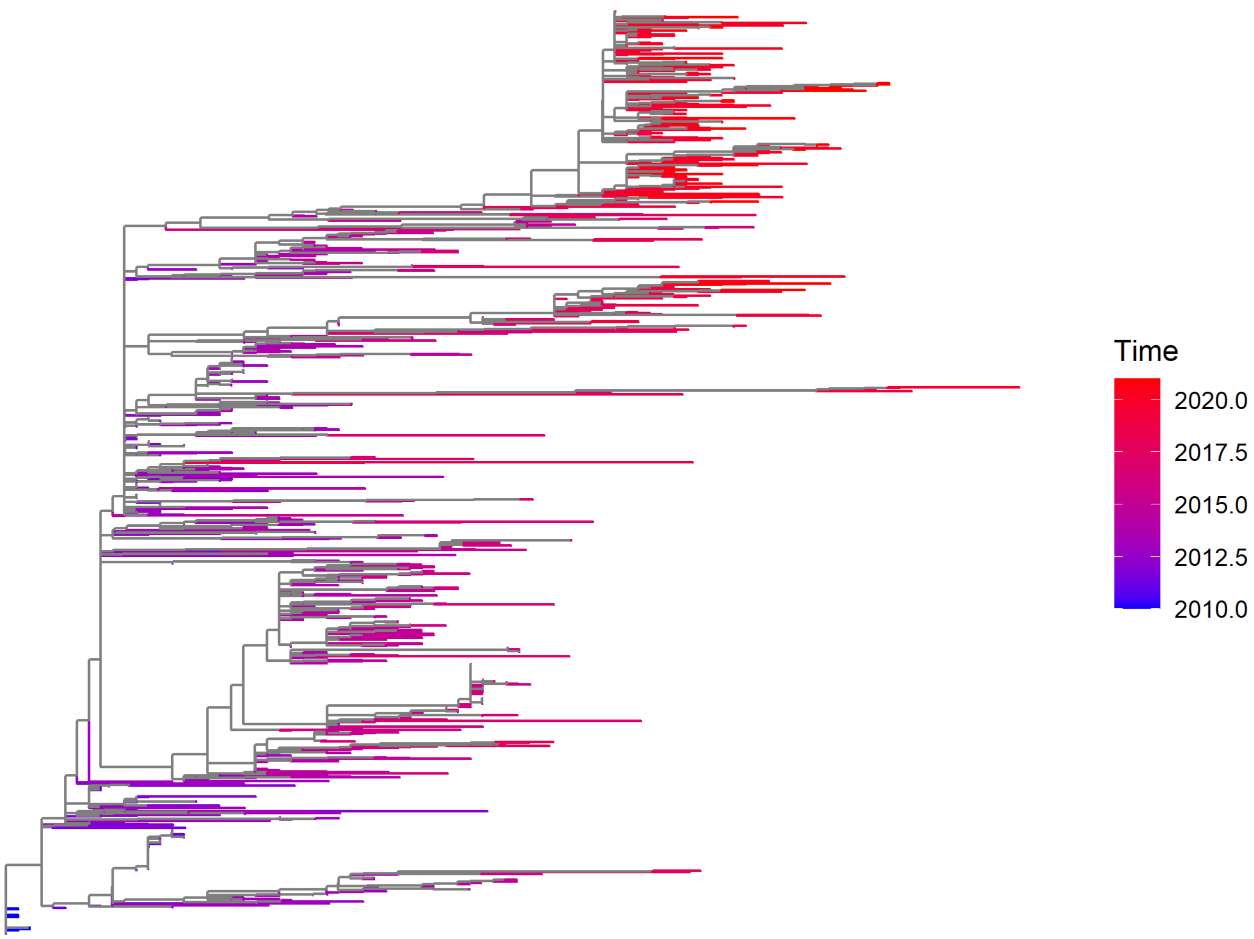


Figure S1. H3 C-IVA maximum likelihood tree. Branches are colored by the progression of time on a blue-to-red gradient beginning in 2010 and ending in March 2021.

**Supplementary Table 1.** Results from a pair of hemagglutination inhibition (HI) assays performed on November 13, 2020 and January 22nd, 2021.

| Antigen name | Passage | A/BEIJING/32/1992 (EXP604 P#32) | A/BEIJING/32/1992 (EXP604P#33) | A/WUHAN/359/1995 (EXP548 P#396) | A/WUHAN/359/1995 (EXP548 P#397) | A/SWINE/NEW YORK/A01104005/2011 (EXP548 P#432) | A/SWINE/NEW YORK/A01104005/2011 (EXP548 P#433) | A/SWINE/IOWA/A01480656/2014 (EXP592 P#721) | A/SWINE/IOWA/A01480656/2014 (EXP592 P#722) | A/SWINE/NEBRASKA/A01567651/2015 (EXPFLU1 P#303) | A/SWINE/NEBRASKA/A01567651/2015 (EXPFLU1 P#304) | A/SWINE/MINNESOTA/A01280592/2013 (EXP569 P#853) | A/SWINE/MINNESOTA/A01280592/2013 (EXP569 P#854) | **A/SWINE/NORTH CAROLINA/A02245294/2019** (FLU41B P#463) | **A/SWINE/NORTH CAROLINA/A02245294/2019** (FLU41B P#464) | A/SWINE/WYOMING/A01444562/2013 (EXP592B P#712) | A/SWINE/WYOMING/A01444562/2013 (EXP592B P#712) | A/SWINE/INDIANA/A01202866/2011 (EXP548 P#414) | A/SWINE/INDIANA/A01202866/2011 (EXP548 P#415) | Antigenic Motif | H3 Clade |
|---|---|---|---|---|---|---|---|---|---|---|---|---|---|---|---|---|---|---|---|---|---|
| A/BEIJING/32/1992 | MDCK2 | **320** | **640** | 160 | 160 | 80 | 80 | 160 | 160 | 20 | 80 | 80 | 80 | * | * | * | * | * | * | NHKEYR | HuVac |
| A/BEIJING/32/1992 | MDCK2 | * | * | * | * | 80 | 80 | 40 | 40 | * | * | * | * | 40 | 40 | 10 | 40 | 40 | 40 | | |
| A/WUHAN/359/1995 | MDCK2 | 40 | 80 | **320** | **640** | 80 | 80 | 320 | 640 | 80 | 80 | 80 | 80 | * | * | * | * | * | * | KHKEYS | HuVac |
| A/WUHAN/359/1995 | MDCK2 | * | * | * | * | 40 | 40 | 160 | 320 | * | * | * | * | 320 | 160 | 80 | 640 | 80 | 160 | | |
| A/SWINE/NEW YORK/A01104005/2011 | MDCK4 | 20 | 80 | 40 | 40 | **1280** | **1280** | 320 | 320 | 80 | 80 | 1280 | 640 | * | * | * | * | * | * | NYNNYK | C-IVA |
| A/SWINE/NEW YORK/A01104005/2011 | MDCK4 | | | | | **640** | **1280** | 160 | 80 | * | * | * | * | 320 | 1280 | 160 | 160 | 160 | 320 | | |
| A/SWINE/IOWA/A01480656/2014 | MDCK3 | 10 | 40 | 160 | 320 | 640 | 640 | **1280** | **2560** | 40 | 20 | 640 | 320 | * | * | * | * | * | * | KYNNYK | C-IVA |
| A/SWINE/IOWA/A01480656/2014 | MDCK3 | * | * | * | * | 320 | 320 | **640** | **1280** | * | * | * | * | 160 | 320 | 640 | 2560 | 160 | 160 | | |
| A/SWINE/NEBRASKA/A01567651/2015 | MDCK2 | 20 | 160 | 40 | 40 | 640 | 320 | 320 | 160 | **2560** | **1280** | 320 | 160 | | | | | | | SYKNYK | C-IVA |
| A/SWINE/NEBRASKA/A01567651/2015 | MDCK2 | | | | | 320 | 320 | 640 | 640 | * | * | * | * | 640 | 80 | 640 | 640 | 40 | 80 | | |
| A/SWINE/MINNESOTA/A01280592/2013 | MDCK2 | 20 | 40 | 40 | 40 | 640 | 640 | 80 | 160 | 160 | 80 | **2560** | **2560** | * | * | * | * | * | * | NYHNYK | C-IVB |
| **A/SWINE/NORTH CAROLINA/A02245294/2019** | MDCK2 | 20 | 80 | 40 | 40 | 640 | 320 | 160 | 160 | 80 | 40 | 1280 | 640 | * | * | * | * | * | * | NYHNYK | C-IVA |
| **A/SWINE/NORTH CAROLINA/A02245294/2019** | MDCK2 | * | * | * | * | 320 | 320 | 160 | 160 | * | * | * | * | **640** | **1280** | 80 | 160 | 80 | 160 | | |
| A/SWINE/WYOMING/A01444562/2013 | MDCK2 | * | * | * | * | 320 | 320 | 640 | 640 | * | * | * | * | 160 | 320 | **1280** | **5120** | 1280 | 640 | KYNNYK | C-IVA |
| A/SWINE/INDIANA/A01202866/2011 | MDCK3 | * | * | * | * | 40 | 80 | 80 | 40 | * | * | * | * | 40 | 80 | 10 | <10 | **2560** | **2560** | NYHGHE | C-IVC |
| **A/SWINE/OKLAHOMA/A01770191/2015** | MDCK2 | 20 | 80 | 20 | 40 | 640 | 1280 | 80 | 80 | 160 | 40 | 1280 | 640 | * | * | * | * | * | * | NYNNYK | C-IVA |
| **A/SWINE/OKLAHOMA/A01770191/2015** | MDCK2 | * | * | * | * | 320 | 640 | 160 | 80 | * | * | * | * | 320 | 640 | 80 | 320 | 80 | 40 | | |
| **A/SWINE/MINNESOTA/A02266068/2018** | MDCK2 | 20 | 40 | 20 | 20 | 640 | 640 | 160 | 160 | 80 | 40 | 640 | 640 | * | * | * | * | * | * | NYNNYK | C-IVA |
| **A/SWINE/MINNESOTA/A02266068/2018** | MDCK2 | * | * | * | * | 640 | 640 | 160 | 160 | * | * | * | * | 160 | 640 | 80 | 160 | 80 | 80 | | |

| | |
|---|---|
| (shaded) | Homologous HI titer |
| * | Not tested (Merged Table) |

**Supplementary Table 2.** Results from Neuraminidase Inhibition (NI) Assay.

| | NA Lineage | 02A.1 | | 02A.2 | | 02B.1 | | 02B.2 | |
|---|---|---|---|---|---|---|---|---|---|
| | | A/swine/Nebraska/ A01492366/2014 (FLU17 P#720) | A/swine/Nebraska/ A01492366/2014 (FLU17 P#721) | A/swine/New York/A01104005/2011 (FLU17 P#722) | A/swine/New York/A01104005/2011 (FLU17 P#723) | A/swine/Minnesota/ A01678475/2016 (FLU23 P#513) | A/swine/Minnesota/ A01678475/2016 (FLU23 P#514) | A/swine/Iowa/ A01480656/2014 (FLU17 P#724) | A/swine/Iowa/ A01480656/2014 (FLU17 P#725) |
| **A/swine/Nebraska/ A01492366/2014** | **02A.1** | **320** | **320** | 160 | 40 | 80 | 20 | 40 | 20 |
| **A/swine/New York/A01104005/2011** | **02A.2** | 160 | 80 | **640** | **160** | 160 | 40 | 40 | <10 |
| **A/swine/Minnesota/A02266068/2018** | **02A.2** | 160 | 160 | 640 | 320 | 320 | 160 | 160 | 10 |
| **A/swine/Minnesota/ A01678475/2016** | **02B.1** | 20 | 40 | 160 | 80 | **640** | **160** | 40 | <10 |
| **A/swine/Iowa/ A01480656/2014** | **02B.2** | 160 | 160 | 320 | 80 | 80 | 80 | **5120** | **640** |
| **A/swine/Oklahoma/A01770190/2015** | **02B.2** | 320 | 80 | 320 | 40 | 160 | 80 | 2560 | 640 |
| **A/swine/North Carolina/A02245294/2019** | **02B.2** | 320 | 160 | 320 | 80 | 160 | 80 | 2560 | 640 |
